# Histological assessment of integrated human cortical organoid grafts after controlled cortical impact

**DOI:** 10.64898/2026.07.20.739683

**Authors:** Caitlyn Smith, Sarah Hamimi, Mackenzie Castellanos, Emma Noel, Ana Paulina Serrano, Said Inaltekin, Nirvaa Shah, Willam Perez, Frank J. Rauscher, Jaeha Kim, Hailong Song, Victoria E. Johnson, Han-Chiao Isaac Chen, Dennis Jgamadze

## Abstract

Rodent models are a mainstay of traumatic brain injury (TBI) research, including investigations into the pathophysiology and treatment of this condition. However, there are fundamental molecular and cellular differences between rodent and human neurons, as well as other cells of the brain. Brain organoids derived from human pluripotent stem cells recapitulate key features of the human brain and have been used to model a variety of neurological disorders. Here, we developed a novel *in vivo* model of human TBI based on controlled cortical impact (CCI) injuries of human organoid grafts transplanted into the brains of young adult rats. Cortical organoids derived from human induced pluripotent stem cells (iPSCs) were grown for 50-60 days *in vitro* before transplantation into rat visual cortex. Injures were performed 2 months later, and histological outcomes were examined at 7 or 30 days after injury. Injury cavities in the integrated grafts were identified at both endpoints with a progression toward larger cavities sizes with time. The injured human tissue exhibited evidence of neuroinflammation with elevated numbers of IBA1+ cells and axonal injury with APP+ cells. There was evidence of increased cell proliferation in the injured grafts acutely after injury that decreased with time. The injured grafts also showed evidence of phosphorylated tau aggregates and accumulation of PNAG, a polysaccharide associated with microbial pathogens. These results support the feasibility of using human organoid grafts in rats as a model of TBI, potentially including the study of long-term neurodegeneration and microbial penetration of the brain after injury.

## Introduction

Traumatic brain injury (TBI) is a major cause of death and long-term neurological disability in the United States.^1^ In addition to the direct mechanical impact of the primary injury, TBI initiates evolving secondary processes that include neuroinflammation, excitotoxicity, oxidative stress, vascular disruption, and cell death.^2,3^ Experimental animal models, especially those based on rodents, have been essential for defining these mechanisms and relating impact parameters to tissue pathology.^4,5^ However, the human brain differs from rodents brains in a number of critical ways, including size and anatomical organization. Differences in developmental programs, cellular processes, molecular pathways, and network properties also exist.^6–9^ These species differences may explain why therapeutics that effectively treat TBI in rodents have failed to translate to humans and motivate the search for complementary models that enable the study of traumatic injury in human neural tissue under controlled conditions.

Brain organoids derived from human pluripotent stem cells reproduce key features of the human brain, including human-specific neural progenitor populations such as outer radial glial, cortical fate specification, and the formation of cortical lamina.^10–12^ These similarities to the human brain have inspired the use of brain organoids as models for a variety of neurological disorders and conditions.^13,14^ More recently, brain organoids have also been explored as models for TBI.^15^ These efforts have thus far generated *in vitro* organoid injury platforms, which have demonstrated trauma-associated neuronal loss, glial reactivity, and tau- and TDP-43-related proteinopathies.^16–18^ Because of their *in vitro* nature and the limited cell populations found within standard organoids, these models do not take into account inflammatory and vascular changes after TBI. Furthermore, there are biomechanical differences in the transmission of forces between neural tissue *in vitro* and brain tissue constrained by the skull and other related tissues.

Transplantation of brain organoids into rodent cortex could address some of the limitations of *in vitro* organoid models of TBI, providing organoids with the environment of the living brain. The feasibility of brain organoid transplantation has been established. ^19–21^ Organoid grafts survive robustly after transplantation with rapid vascularization by the host animal.^20^ Neurons within the grafts mature over time and integrate with the host brain both structurally via synaptic connectivity and functionally into brain networks.^19,21,22^

Here, we developed and characterized an *in vivo* model of human TBI based on transplanted organoids, building on the foundation established by prior *in vitro* organoid TBI models and organoid transplantation studies. Human cortical organoids derived from induced pluripotent stem cells (iPSCs) were allowed to integrate with the host brain for 2 months before being subjected to a standard controlled cortical impact (CCI) injury.^23^ At 7 or 30 days after CCI injury, immunohistochemical analysis of the organoid grafts and surrounding tissue compartments was performed. We observed progressive expansion of cavities within the injured organoid grafts associated with neuronal cell death, glial reactivity, and neuroinflammation. We also found tau aggregation within the injured grafts as well as evidence of microbial penetration after injury.

## Methods

### Maintenance of human induced pluripotent stem cells

The C1.2 human induced pluripotent stem cell (iPSC) line was maintained as feeder-free colonies on hESC-Qualified Matrigel-coated plates (Corning) in mTeSR Plus medium (StemCell Technologies). To coat plates, matrigel was thawed on ice for 1 hr and diluted in cold DMEM/F12 Media (Gibco) at 1% (v/v). Plates were incubated for 2 hrs prior to plating cells. Cultures were passaged approximately 1:10 every 4-6 days using ReLeSR (StemCell Technologies), The iPSC lines were kept below passage 50 to minimize the accumulation of mutations and were PCR tested monthly for mycoplasma using Universal Mycoplasma Detection Kit (ATCC).

### Generation of dorsal forebrain organoids

Dorsal forebrain organoids were generated using a previously published protocol.^10^ In brief, embryoid bodies (EBs) were created by detaching stem cell colonies using 1mL ReLeSR. On Day 0, detached iPSC cells were seeded in ultra-low attachment 96-well U-bottom plates (Corning Costar) in mTeSR plus media and 10uM ROCK inhibitor Y-27632 (StemCell Technologies). From day 1 to 5, resuspended EBs were cultured in ultra-low-attachment 6-well plates (Corning Costar) in DA induction media consisting of DMEM/F12 media, 20% KSR, 2 mM GlutaMAX, 0.1 mM non-essential amino acids (NEAA), 94.6 µM β-mercaptoethanol, 200 nM LDN193189 (Tocris), 10 µM SB431542 (Tocris) and 100 U/mL penicillin and 100 ng/mL streptomycin. On day 6, the media was half-changed to CS induction media (IM) consisting of DMEM/F12 with 1X N2 supplement (Gibco), 0.1 mM NEAA, 2 mM GlutaMAX, 1 µM CHIR 99021 (Tocris), 1 µM SB431542 (Tocris), and 100 U/mL penicillin and 100 ng/mL streptomycin. On day 7, EBs were embedded in a 1:1 Matrigel (Corning) to IM mixture. This mixture was maintained on ice, and EBs were dispersed throughout it by gentle pipetting. The Matrigel and embedded EBS were then spread on an ultra-low attachment 6-well plate and allowed to gel at 37°C for 60 min. After 60 minutes, 3mL IM mixture was gently added to each well. The embedded EBs were then cultured in stationary conditions in IM media from day 7-14. On day 14, the embedded EBs were gently broken out of the Matrigel hydrogel with a 5-mL serological pipette and transferred to an orbital shaker (100-120 rpm). From day 14 to 71, the organoids were cultured in differentiation media consisting of 50% DMEM/F12 and 50% Neurobasal medium (Gibco) with 1X N2 supplement, 1X B27 supplement (Gibco), 2 mM GlutaMAX, 2.8 ng/ml human insulin (Sigma), 0.1 mM NEAA, 100 μM β-mercaptoethanol, and 100 U/mL penicillin and 100 ng/mL streptomycin. Media changes were performed every 2 days. Thereafter, the organoids were maintained in maturation media consisting of Neurobasal with 1X B27 supplement, 2 mM GlutaMax, 0.2 mM ascorbic acid, 20 ng/mL brain-derived neurotrophic factor (PeproTech), 20 ng/mL glial cell line-derived neurotrophic factor (PeproTech), and 50 U/mL penicillin and 50 ng/mL streptomycin. Media changes were performed every 2 days. Organoids were sliced to 500 µm using a vibratome (Leica Biosystems) on day 40.

Only organoids that passed strict quality control were selected for subsequent transplantation. Metrics that were assessed including the clearing of embryoid body borders before Matrigel embedding suggestive of the formation of radially organized neuroepithelium and the development of defined buds in Matrigel without cyst formation or evidence of premature differentiation.

### Animal procedures and institutional approvals

All animal experiments described in this study were approved by the Institutional Animal Care and Use Committee (IACUC) at the University of Pennsylvania (protocol number 805600) and were conducted in accordance with the National Institutes of Health’s Guide for the Care and Use of Laboratory Animals. Animals were housed in pairs under standard conditions and kept on a 12-hr light/dark cycle (lights on at 6:00 a.m.) with *ad libitum* access to food and water.

### Study design

The experimental sequence consisted of organoid transplantation, CCI injury to the graft region 2 months later, and then brain tissue harvesting 7 or 30 days after CCI. Experimental groups included CCI injury only and organoid transplant plus CCI, each with independent 7- and 30-day endpoints.

### Organoid transplantation

Young adult Long Evans rats (male, 250-300 g, 8-12 weeks) assigned to organoid transplantation groups received intraperitoneal injections of 10 mg/mL Sandimmune cyclosporine A (Novartis Pharmaceuticals Corporation) beginning 2 days before surgery and daily thereafter until animal sacrifice. Anesthesia was induced with 5% isoflurane and maintained at 2.5%. Animals were positioned in a stereotaxic frame, and body temperature was supported with a water circulation-based heating pad (Gaymar). Depth of anesthesia was monitored by observation of the animals’ respiration and toe pinch responses. Prior to skin incision, animals were given dexamethasone (IP, 1 mg/kg; Mylan) and bupivacaine (subcutaneous, 2 mg/kg; APP Pharmaceuticals). A 5-mm circular craniotomy was made centered at 5 mm posterior to bregma and 2.5 mm lateral to midline. Following durotomy, a cortical cavity was created by vacuum aspiration, and a day 50-60 organoid was manually placed in the cavity with a cut-tip 200uL pipette. Organoids were incubated with 20 µg/mL necrostatin-1 (Enzo Life Sciences) for 24 hours prior to transplantation. The craniotomy was sealed with a custom 1.5-mm thick polydimethylsiloxane (PDMS; Dow Silicones Corporation) cap covered with RelyX™ Unicem 2 Automix Self-Adhesive Resin Cement (3M) and the incision was closed with suture. Meloxicam SR (subcutaneous, 2mg/mL, Wedgewood) was administered postoperatively for systemic pain control for at least 3 days after the procedure.

### Controlled cortical impact injury

After induction of anesthesia and positioning in a stereotaxic frame, the host brain or region of the organoid graft was exposed. For the CCI optimization study, impact depths of 1, 2, or 3 mm were tested with a 3-mm-diamter impactor tip with a peak velocity of 2.5 m/s. Subsequently, studies were performed using a impact depth of 2 mm. Following CCI injury, a new cranioplasty (PDMS cap secured in place with bone cement) was fashioned, and the skin was closed with suture.

### Frozen section immunohistochemistry

Animals were euthanized with Euthasol (intraperitoneal, >150 mg/kg) at the designated endpoint, and tissue processing was performed followed our established protocols for frozen immunohistochemistry.^19^ In brief, cyrosectioned samples were blocked with 5% normal goat serum (Vector Labs) and 0.3% Triton X-100 (Sigma), incubated with primary antibodies overnight at 4°C, rinsed with tris-buffered saline with Tween 20 (TBST), incubated with secondary antibodies for 2 hours at room temperature, rinsed with TBST, and then coverslipped with Fluoromount-G (Fisher). For CTIP2 and SATB2 antibodies, antigen retrieval prior to TBST incubation was necessary. Slides were dried for 4hrs at RT and then placed in a pressure cooker in microwave with boiling 1X Tris-EDTA buffer (BioLengend) for 8 minutes. After cooling for 10 minutes, slides were then incubated in TBST and primary antibodies.

### Paraffin immunohistochemistry

Paraffin immunohistochemistry was performed as previously described^24^. Specifically, 8 μm sections were prepared from formalin-fixed paraffin-embedding tissue blocks. Sections were deparaffinized and rehydrated through exchanges of xylene, 100%, 95% ethanol, and water. Hydrogen peroxide (3%) was used to quench endogenous peroxidase activity for 15 mins, followed by heat-induced antigen retrieval in Tris-EDTA buffer (pH 8.0) or Citrate Buffer (pH 6.0). Next, slides were blocked with normal serum (Vector Labs) in Optimax buffer (BioGenex) for 30 mins at room temperature (RT) and then incubated with primary antibodies overnight at 4°C in a humidified chamber. Primary antibodies used in this study included amyloid precursor protein (APP) (MAB348, Millipore, 1:80,000) and AT8 (MN1020 Thermo Fisher, 1:2,000). Next day, the sections were further incubated with the corresponding biotinylated secondary antibodies (1:250, Vector Labs) followed by visualization using avidin biotin complex (ABC) and 3,3′-Diaminobenzidine (DAB) peroxidase substrate application per the manufacturer’s protocols. Sections were counterstained with hematoxylin.

### Image analysis

Stained slides were scanned on a Hamamatsu NanoZoomer S360 at 20x and analyzed in QuPath v0.6.0. One slide per antibody per animal was selected, with up to six 25-micrometer thick coronal sections represented. In transplant conditions, SC121 immunoreactivity was used to define graft-associated annotations. For nuclear markers, DAPI-based cell detection was performed within the primary graft region, and detections were classified using marker-specific intensity thresholds. For IBA1 and GFAP, positive fluorescent area was recorded within the graft region and within a surrounding 500-micrometer region.

### Statistical analysis

Statistical analyses were performed using the animal as the experimental unit. When multiple histological sections were analyzed from the same animal, section-level measurements were averaged to generate one animal-level mean for each outcome and region of interest. Descriptive statistics were calculated from these animal-level means and are reported as mean ± standard deviation. Comparisons between the independent 7- and 30-day cohorts were performed using two-sided Mann–Whitney U tests. Qualitative endpoints and pilot comparisons not designed for hypothesis testing were summarized descriptively without inferential analysis. Throughout the manuscript, n denotes the number of animals, all reported p values are two-sided, and p < 0.05 was considered statistically significant.

## Results

### Optimization of CCI injury parameters for human organoid grafts

Establishing an *in vivo* organoid injury paradigm first required determining the impact parameters that resulted in significant injury of the graft tissue without eliminating the tissue altogether. The impact velocity was set at 2.5 m/s, and three different impactor depths from the brain surface (1-, 2-, and 3-mm) were compared with histology 7 days after CCI injury of 2-month-old human cortical organoid grafts (Fig. 1A-C).

**Figure 1.**
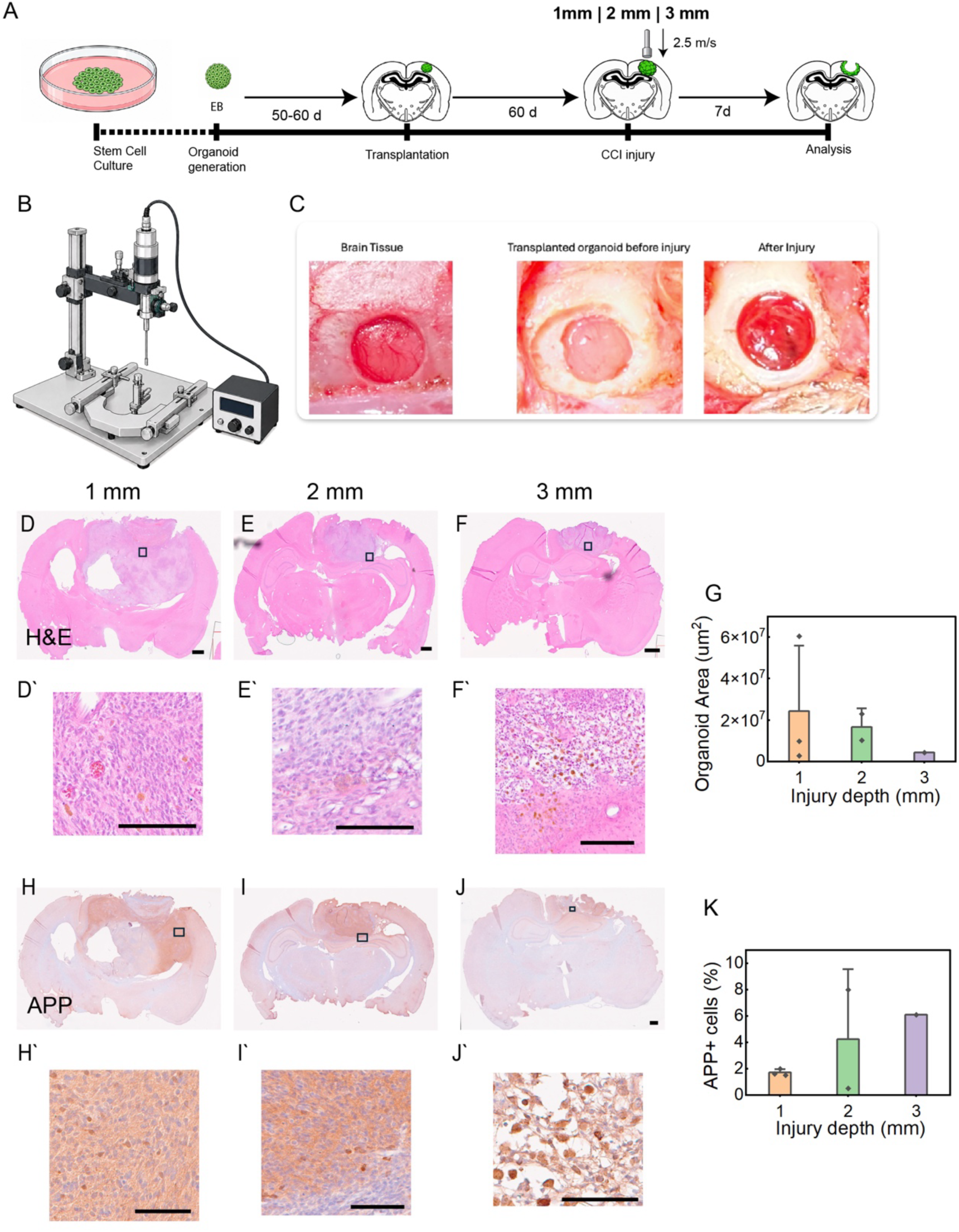
Optimization and establishment of an in vivo organoid-targeted controlled cortical impact paradigm. (A) Experimental timeline. Human cortical organoids were generated for 50-60 days, transplanted into rat cortex, allowed to integrate in vivo for 60 days, and subjected to controlled cortical impact (CCI) at 2.5 m/s using impact depths of 1, 2, or 3 mm. All pilot animals were analyzed 7 days after injury. (B) CCI apparatus used to position and deliver the impact. (C) Representative gross surgical views of exposed brain tissue, the transplanted organoid before injury, and the targeted site after injury. Histological panels are aligned by impact depth throughout: the left column shows 1 mm, the middle column 2 mm, and the right column 3 mm. (D-F) Representative hematoxylin and eosin (H&E)-stained coronal sections from the 1-, 2-, and 3-mm groups, respectively. (D’-F’) Higher-magnification views of the boxed regions in D-F. (G) Cross-sectional organoid area at each impact depth. (H-J) Representative APP immunohistochemistry from the 1-, 2-, and 3-mm groups, respectively. (H’-J’) Higher-magnification views of the boxed regions in H-J. (K) Percentage of detected cells classified as APP-positive within the analyzed organoid regions. Individual observations are shown; bars and error bars summarize the corresponding depth groups. Scale bars: D-F and H-J, 1 mm; D’-F’ and H’-J’, 100 µm.

H&E-stained sections showed that the injured graft region remained identifiable at each tested depth, but the amount of retained organoid tissue differed across the depths (Fig. 1D-G). Cross-sectional area was greatest at an impactor depth of 1 mm, intermediate at 2 mm, and smallest at 3 mm. Amyloid precursor protein (APP) immunoreactivity showed an inverse relationship with the least amount of positive staining at a depth of 1 mm and the greatest at 3 mm (Fig. 1H-K).

These two readouts together revealed the central tradeoff in the model. Injury at a depth of 1 mm left more tissue available for histological analysis but showed comparatively little tissue injury as judged by APP labeling, whereas an injury depth of 3 mm produced the greatest amount of APP labeling with little retained organoid area.

Injury at a depth of 2 mm occupied an intermediate window, providing easily detectable injury-associated APP labeling while preserving a significant amount of graft tissue for histological analysis. Thus, 2 mm was chosen as the injury depth for the subsequent experiments.

### Injury outcomes at 7 and 30 days after CCI

We next asked whether the selected 2-mm impact depth generated a reproducible lesion with evidence of tissue and cellular damage. Cavities were evident in both injured rat cortex and brains bearing injured organoid grafts at 7 and 30 days after CCI (Fig. 2B-E, F). For both injury contexts, mean cavity area progressively increased from 7 to 30 days after injury, although this result did not reach statistical significance. The cavity size in injured rat cortex was 5.77 ± 2.58 × 10⁶ at 7 days and 13.45 ± 11.24 × 10⁶ µm² at 30 days (7d N=3, 30d N=2, P=0.508). The mean cross-sectional area in the injured grafts was 4.4 ± 2.1 × 10⁶ µm² at 7 days and 10.0 ± 6.5 × 10⁶ µm² at 30 days (N=4, p=0.11235). As indicated by the large standard deviation at 30 days, we observed substantial heterogeneity in the size of the injury cavities within the organoid grafts.

**Figure 2.**
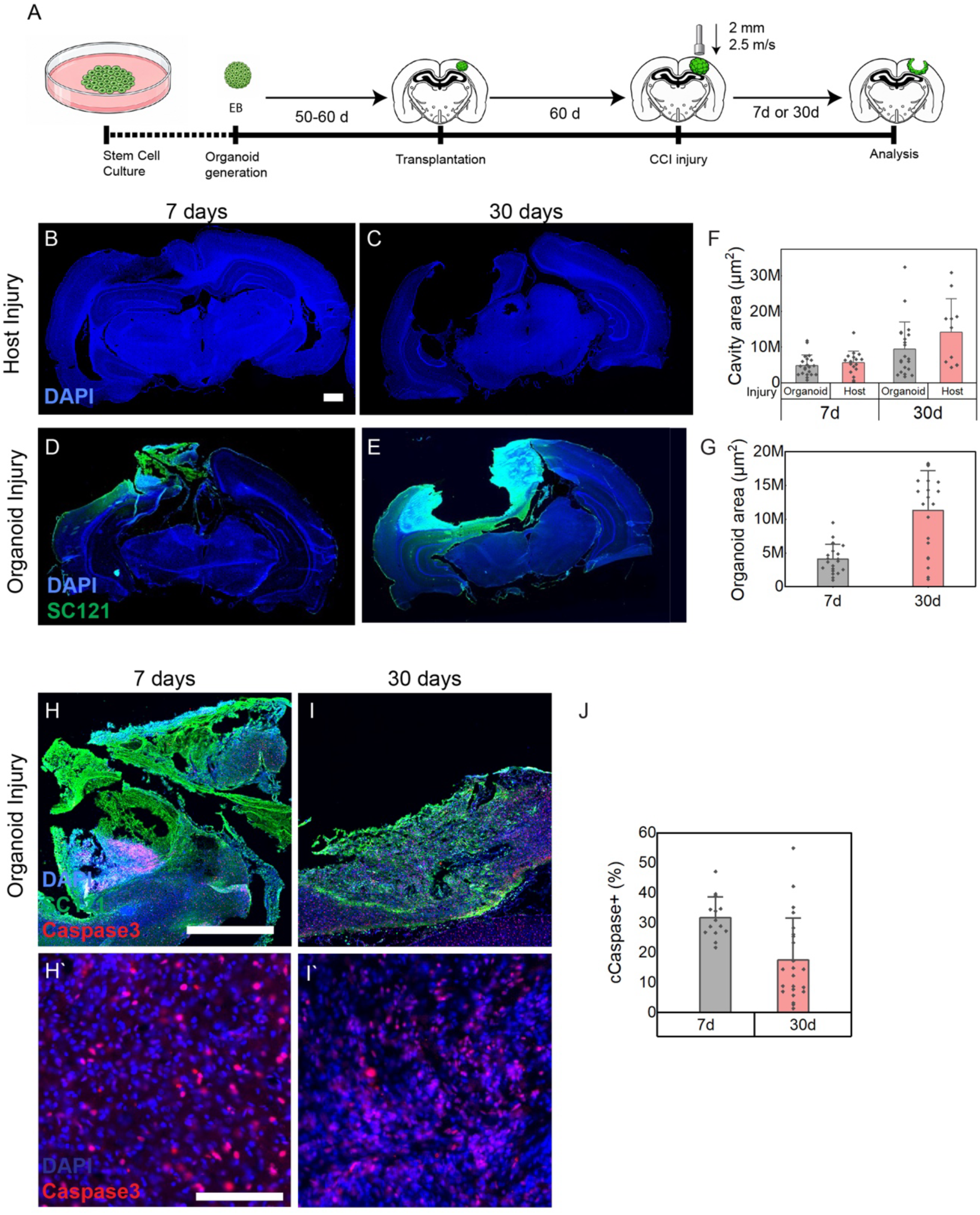
Organoid graft morphology and cleaved caspase-3 labeling after controlled cortical impact. (A) Experimental workflow. Human cortical organoids were generated for 50-60 days, transplanted into rat cortex, allowed to integrate in vivo for 60 days, and subjected to CCI at 2-mm depth and 2.5 m/s. Independent cohorts were analyzed 7 or 30 days after CCI. (B-C) Representative whole-section images from host injury-only animals at 7 and 30 days, respectively; nuclei are labeled with DAPI (blue). (D-E) Representative whole-section images from organoid injury animals at 7 and 30 days, respectively; DAPI is shown in blue and SC121-positive graft-associated human tissue in green. (F) Injury cavity area in organoid targeted (Organoid) and host injury only (Host) sections at each endpoint. (G) Cross-sectional organoid-region area in analyzed 7- and 30-day sections. (H-I) Representative graft-region images at 7 and 30 days showing DAPI (blue), SC121 (green), and cleaved caspase-3 (red). (H’-I’) Higher-magnification views showing DAPI and cleaved caspase-3 within the graft-region annotation. (J) Percentage of DAPI-detected cells classified as cleaved caspase-3-positive within the graft-region region of interest (ROI). Each point in F, G, and J represents one analyzed section nested within an animal; bars show mean ± SD. Scale bars: B-E, 1 mm; H-I, 500 µm and H’-I’, 100 µm.

Organoid grafts were confirmed to be of human origin using the human-specific marker SC121 (Fig. 2D-G). Within the graft region, nuclei positive for cleaved caspase 3 were detected at both endpoints (Fig. 2H-J). The positive fraction averaged 31.7 ± 7.0% at 7 days (14 sections from 3 animals) and 17.6 ± 14.0% at 30 days (23 sections from 4 animals, P=0.37676). The injured organoid grafts contained more apoptotic cells than has been reported for uninjured organoid grafts,^19^ although no change in the frequency of apoptotic cells could be detected between time points. Together, these structural and cellular metrics demonstrated that the selected CCI parameters produced a substantial lesion with evidence of significant cell death.

### Neuronal compartments within the injured organoid grafts

Next, we evaluated the state of neural progenitors and neurons within the organoid grafts. Significant numbers of PAX6+ neural progenitors were observed at both 7 and 30 days after injury, as expected based on the early developmental status of the organoids (Fig. 3F-H). We specifically examined proliferating cells using Ki67 staining. Ki67-positive cells were present within the graft region at both endpoints with a trend toward more proliferating cells at 7 days (Fig. 3A-E). The Ki67-positive fraction averaged 33.8 ± 16.0% at 7 days and 10.5 ± 7.3% at 30 days (16 and 23 sections, respectively, from 4 animals per endpoint, P=0.0606).

**Figure 3.**
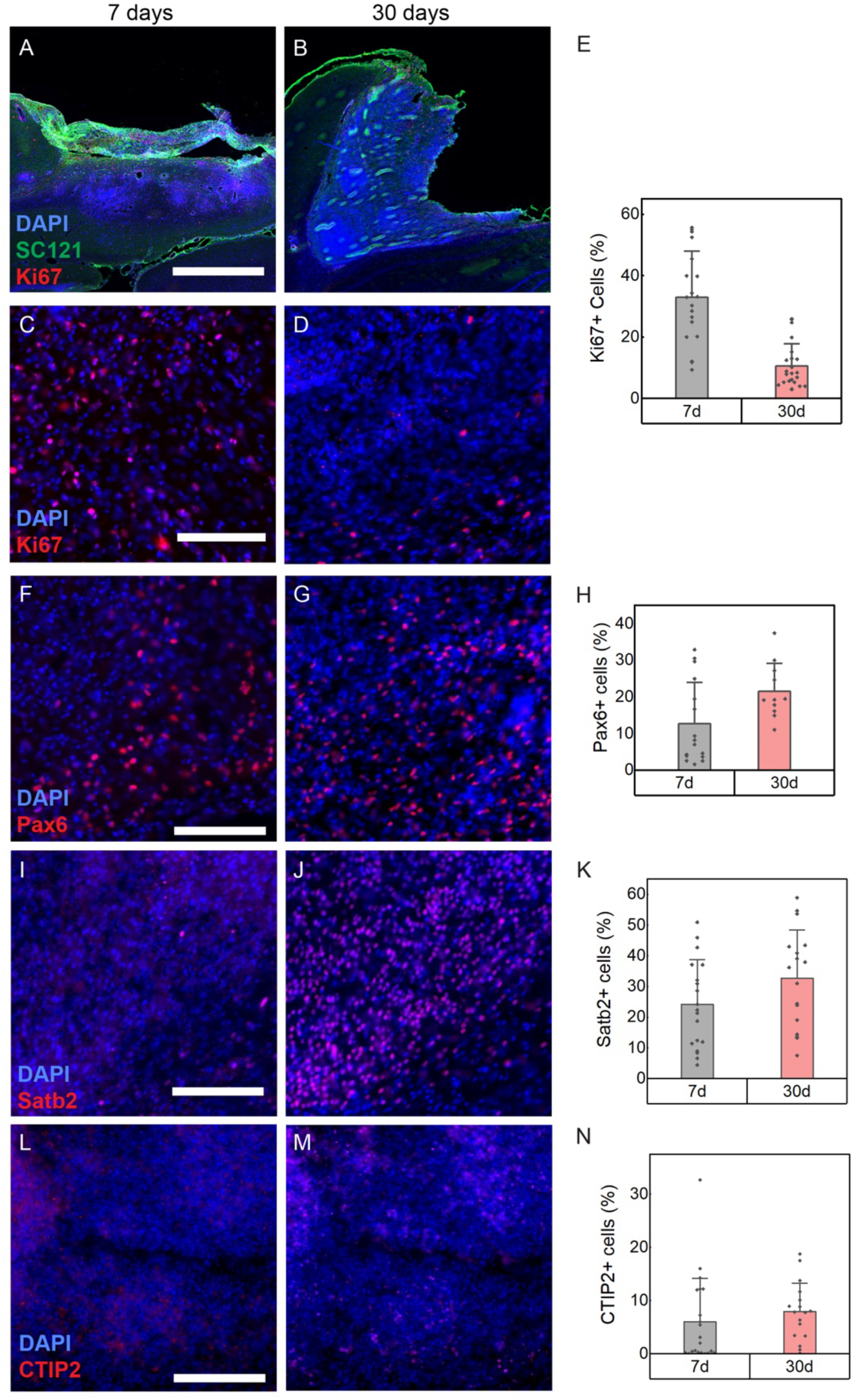
Cortical organoid cellular phenotype after organoid-targeted controlled cortical impact. (A-B) Representative low-magnification fluorescence images of graft-associated tissue collected 7 or 30 days after CCI. Nuclei are labeled with DAPI (blue), SC121 immunoreactivity identifies graft-associated human tissue (green), and Ki67 immunoreactivity is shown in red. (C-D) Higher-magnification images of DAPI and Ki67 labeling. (E) Ki67-positive nuclei expressed as a percentage of total detected nuclei within the graft-region annotation. (F-G) Representative DAPI and PAX6 labeling at 7 and 30 days, with corresponding quantification in H. (I-J) Representative DAPI and SATB2 cells, with corresponding quantification in K. (L-M) Representative DAPI andCTIP2 labeling, with corresponding quantification in N. Scale bars: A-B, 1 mm; C-M 100 µm.

We also assessed cortical layer-specific neurons using the upper-layer marker SATB2 and lower-layer marker CTIP2. In both cases, similar numbers of positive cells were identified at both endpoints. For SATB2, there were 24.1 ± 14.6% at 7 days and 32.6 ± 15.9% at 30 days (N=4, P=0.66501). For CTIP2, there were 6.0 ± 8.2% at 7 days and 7.9 ± 5.4% at 30 days (N=4, P=0.47049) (Fig. 3I-N). These data suggest that traumatic injury did not preferentially affect upper-versus lower-layer cortical neurons.

### Graft-associated glial reactivity and neuroinflammation

Because the grafts were injured within a living host brain, we asked whether there was evidence of glial reactivity and inflammation within the organoid grafts. GFAP immunoreactivity was present at low levels in both the grafts as well as the adjacent host brain (Fig. 4G-L). GFAP-positive area within the graft region averaged 9.5 ± 5.9% at 7 days and 11.3 ± 8.4% at 30 days (N=4, P=1).; in the surrounding border, the corresponding values were 1.76 ± 1.19% and 3.26 ± 2.42% (N=4, P=0.64343). IBA1 immunoreactivity was detected in both compartments at both endpoints (Fig. 4A-F).

**Figure 4.**
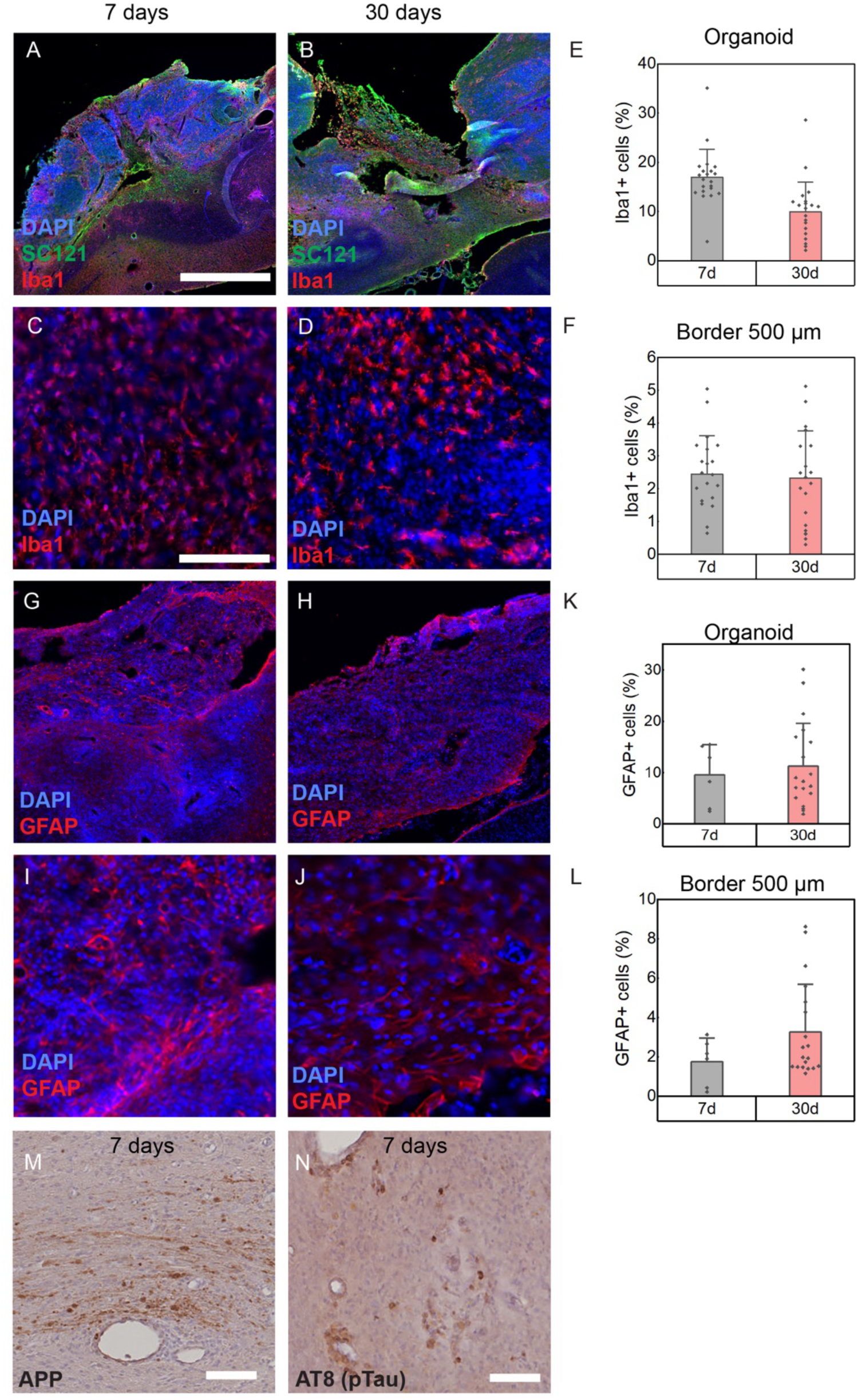
Cortical organoid and border host tissue resolved inflammation and injury markers after organoid-targeted controlled cortical impact. (A-D) Representative fluorescence images at 7 days (A, C) and 30 days (B, D) after CCI. Low-magnification images (A, B) show DAPI-labeled nuclei (blue), SC121-positive graft-associated human tissue (green), and IBA1 immunoreactivity (red); higher-magnification graft-region fields (C, D) show DAPI and IBA1 labeling. (E-F) IBA1-positive fluorescent area expressed as a percentage of the graft-region area (E) or the surrounding 500-µm border area (F). (G-J) Representative DAPI (blue) and GFAP (red) fluorescence at 7 days (G, I) and 30 days (H, J), shown at lower (G, H) and higher (I, J) magnification. (K-L) GFAP-positive fluorescent area expressed as a percentage of the graft-region area (K) or the surrounding 500-µm border area (L). (M-N) Representative brightfield APP (M) and tau (N) immunohistochemistry from 7-day post-CCI samples. These panels are included for qualitative, descriptive purposes only and were not quantified. Each point in E-F and K-L represents one analyzed section nested within an ani mal, and bars show mean ± SD. Scale bars: A-B, 1 mm; C-J 100 µm. M-N 500 µm.

Within the graft region, IBA1 positive averaged 17.0 ± 5.7% at 7 days and 9.9 ± 6.1% at 30 days (N=4, P=0.11235). In a 500 µm band of host cortex directly adjacent to the organoid grafts, the corresponding values were nearly unchanged at 2.45 ± 1.17% and 2.32 ± 1.44% (N=4, P=1). Thus, IBA1 labeling was concentrated within the graft region rather than the immediately surrounding host tissue. Because neither IBA1 nor GFAP was paired with species-specific markers, it is not clear whether positive cells were of human graft of rat host origin. Representative APP-immunoreactive profiles and AT8 immunoreactivity were also observed in the tissue samples (Fig. 4M, N), indicating the presence of axonal pathology and hyperphosphorylated tau aggregates, respectively. Taken together, these analyses suggest the presence of microglia-associated inflammation, although the status of astrogliosis is less clear.

### PNAG immunoreactivity within the injured organoid graft

Finally, we asked whether the organoid grafts contained evidence of microbial penetration after TBI. At 30 days after injury, poly-N-acetylglucosamine (PNAG) immunoreactivity was readily visible in the injured organoid region, whereas little signal was evident in contralateral cortex (Fig. 5A-D). Quantification of these data showed a 7.5-fold increase in the presence of PNAG in the injured organoid graft (Fig. 5E).

**Figure 5.**
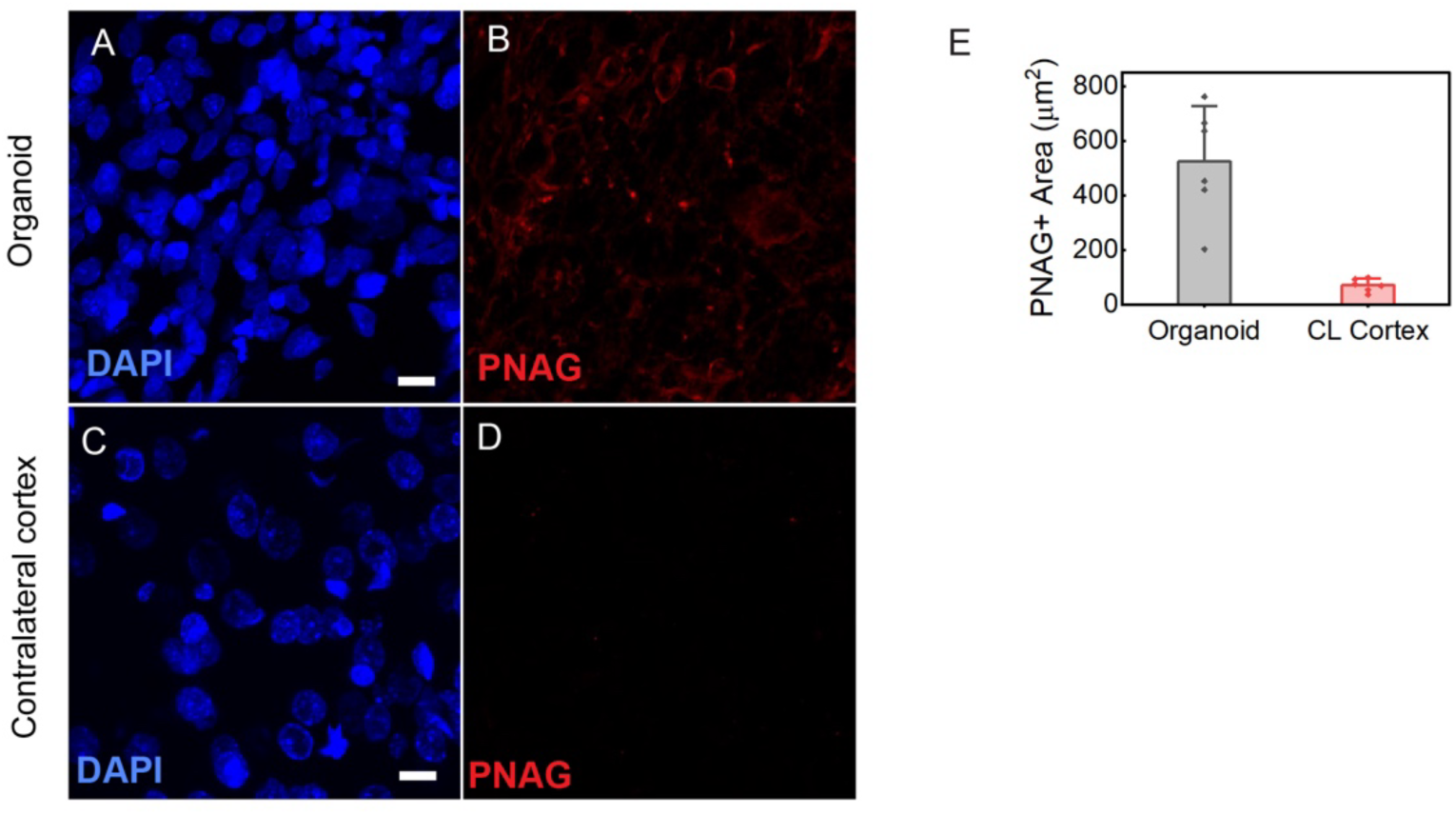
PNAG immunoreactivity in injured organoid tissue and contralateral cortex at 30 days after controlled cortical impact. (A-B) Representative DAPI staining (A; blue) and poly-N-acetylglucosamine (PNAG) immunoreactivity (B; red) in the injured organoid region 30 days after CCI. (C-D) Representative DAPI staining (C; blue) and PNAG immunoreactivity (D; red) in contralateral cortex. (E) Absolute PNAG-positive area in injured organoid samples and contralateral control samples. Each point represents one retained measurement (six per group), and bars show mean ± SD. Scale bars: A D, 10 µm.

## Discussion

Here, we developed a novel experimental model of human TBI based on CCI injuries of 2-month-old human cortical organoids transplanted into rat cortex. We identified CCI parameters that balanced injury severity with the presence of a graft that could be analyzed histologically. Subsequently, we examined the injured human organoid grafts histologically at 7 and 30 days after injury. In addition to expanding cavities, we observed neuronal loss, axonal injury, and cell death within the organoids. There was an increase in reactive astrocytes and microglia, suggestive of an inflammatory microenvironment. Lastly, the grafts contained evidence of tau aggregation and microbial infiltration based on the presence of PNAG. These findings suggest that our model is capable of reproducing key findings of TBI.

### Species differences between rodent and human brains

The anatomic differences between the brains of rodents and humans are obvious. The human brain has a significantly larger surface area than rodent brains, requiring gyrification to fit within the skull. In contrast, the rodent brain is lissencephalic. The increased scale of the human brain necessitates longer axon fibers to facilitate communication between distant brain regions, resulting in a substantially higher white matter-to-grey matter ratio.^25^ The anatomic features of a large, gyrencephalic brain have crucial biomechanical contributions to pathophysiology that cannot be completely captured in rodent models of TBI.^26^

Beyond these gross differences, there are also species differences at the cellular and molecular levels between rodents and humans that may play a role in responses to traumatic injury. Large-scale transcriptomic studies have revealed a significant divergence in RNA transcript expression between mice and human brains.^8^ Some of the greatest differences were seen in ion channels, neurotransmitter receptors, extracellular matrix components, and cell-adhesion molecules,^8,27^ all of which are relevant to TBI. At a cellular level, human neurons exhibit a more compact dendritic architecture^6^ and express a lower density of ion channels compared to rodents,^7^ features that may impart differential metabolic demands and thus cellular vulnerability after injury.

### *In vitro* organoid models of TBI

While human brain organoids do not address the anatomic shortcomings of rodent TBI models, they provide ready access to human neurons and glia with brain-specific architecture. Initial versions of brain organoids used undirected protocols to generate whole brain organoids comprised of neural tissue from multiple brain regions.^11^ More recent organoid protocols have focused on directing differentiation toward a specific brain region in order to obtain greater organoid consistency.^10,12^ Protocols for creating cortical organoids, such as the ones used in this study, promote a dorsal telencephalic fate, resulting in the generation of neural progenitors from the ventricular and subventricular zones. In particular, outer radial glia, which are thought to be responsible for the cortical expansion that make human brains unique from other species, have been identified in cortical organoids.^28^ These organoids develop distinct cortical layers over time,^10^ although full recapitulation of the laminar structure of cortex has not yet been achieved. Glia develops in standard cortical organoids at later time points.^29^ Protocols for accelerated generation of astrocytes^30^ and oligodendrocytes^31^ have also been described.

Several groups have taken advantage of human brain organoids to develop *in vitro* models of TBI. An early report described CCI injuries of organoids placed within a brain phantom composed of agarose and gelatin and mouse skull to mimic the *in vivo* environment.^16^ The organoids were examined 7 days after injury and exhibited evidence of neuronal cell death and astrogliosis. Organoids have also been subjected to uniaxial compression loading.^17^ Transcriptomic analysis showed gene enrichment in the domains of necrotic cell death, nicotinamide adenine dinucleotide synthesis, and interleukin-2 regulation. High-intensity ultrasound has been utilized to induce mechanical injuries in organoids with resultant cell death, tau phosphorylation, and TDP-43 nuclear egress.^18^ A CRISPR interference screen identified KCNJ2, a mechanosensory channel, as a potential therapeutic target to mitigate neuronal death after mechanical injury. Lastly, platforms for studying the effects of blast injury on brain organoids have been developed.^32,33^ These examples highlight the opportunities afforded by human brain organoids to investigate the pathophysiology of TBI in the context of human neural tissue and to screen for and test therapeutics. However, the lack of inflammatory and immune cells, a vascular system, and physical environment of the brain are important limitations.

### CCI injury of human organoid grafts

Our novel *in vivo* model of human TBI yielded several findings that are consistent with standard rodent CCI models. A reproducible cortical cavity formed in the injured organoid graft but not in the adjacent host brain. Injury cavities are a consistent finding of rodent models of CCI.^23,34^ These cavities expand in size over time,^35^ and we observed a trend in this direction in the injured organoid grafts from 7 to 30 days after injury. Other prominent histological features in the injured grafts included a large number of apoptotic cells and microglia infiltration from the host that showed a trend toward greater prominent at 7 days after injury compared to 30 days after injury. The time course of microglia infiltration is similar to what has previously been reported for rodent TBI models.^35^ Interestingly, we did not see significant changes in astrocytes within the injured organoids. The absence of an astrocytic reaction could be explained in part by the small number of astrocytes within transplanted organoids at baseline.^19^ Overall, we observed several histological findings within CCI-injured human cortical organoid grafts that replicated the findings of rodent CCI models.

Traumatic brain injury has long been linked to neurodegenerative diseases such as Alzheimer’s disease and chronic traumatic encephalopathy.^36,37^ Animal models have extensively explored the link between TBI and pathologies related to chronic neurodegeneration, such as aggregates of hyperphosphorylated tau.^38–40^ However, differences exist in rodent versus human tau proteins in terms of the isoforms that exist and their N-terminal sequences, which may have implications for binding patterns with other proteins.^41^ In this study, we observed tau aggregates in the injured organoid grafts, confirming the feasibility of using this model to evaluate the mechanisms of TBI-induced tauopathy and its long-term consequences in human neural tissue. This approach could complement previously established transgenic rodent models expressing human tau.^42^

Poly-β-(1→6)-*N*-acetylglucosamine (PNAG) is an exopolysaccharide produced by many bacterial and fungal pathogens.^43^ Vaccination against PNAG has been explored as a treatment for bacterial meningitis. It has been postulated that PNAG entry into the brain induces chronic neuroinflammation and possibly neurodegenerative diseases. Our initial results support the idea that TBI promotes PNAG entry into the brain, perhaps secondary to breakdown of the blood-brain barrier. Our model is primed for additional studies to examine the mechanism of PNAG entry into the brain and its long-term consequences.

### Limitations of the current model

There are several limitations to the current model that deserve mention, including anatomic, immunological, and cellular factors. While it allows for investigation of the effects of traumatic injury on human neural tissue *in vivo*, the model does not replicate the size or anatomic organization of the human brain. These considerations were the reason why the development of large-animal models of TBI were pursued, such as those in pigs.^26,44^ One potential next step could be the creation of a porcine TBI model based on organoid transplantation. Another strategy could be the transplantation of organoids with cortical folds that mimic the gyrencephlic nature of the human brain. Deletion of PTEN has been shown to induce folding in cortical organoids.^45^

To accommodate human organoid xenografts, host animals in this study were chemically immunosuppressed with cyclosporine A, which raises two issues. Cyclosporine is known to have neuroprotective effects after TBI,^46,47^ which introduces a confounding factor for the results of this study. The immunosuppressant effects of cyclosporine also preclude a comprehensive evaluation of the inflammatory and immune responses after injury in the organoid grafts. The current instantiation of our model does provide some insight into the inflammatory cascade after traumatic injury in human neural tissue, especially when compared to available *in vitro* organoid TBI models. However, future iterations should consider alternative immunological approaches that do not require chemical immunosuppression, such as hosts with humanized immune systems^48^ and hypoimmunogenic cell sources.^49^

Lastly, cellular factors may limit the generalizability of our model. Brain organoids, even after months of *in vitro* or *in vitro* growth, are still relatively immature structures that reflect prenatal and early postnatal human neurodevelopment.^21,28,50,51^ Thus, any model based on stem cell-derived brain organoids may best represent early pediatric TBI rather than adult TBI. It should also be kept in mind that transplantation of currently available brain organoids results in grafts that do not contain all the normal cell types or the normal numbers of glia in the brain,^19^ which may alter responses to traumatic injury.

**Table 1.** Key Resources.

| **Antibody** | **Source** | **Identifier / Cat. #** |
| --- | --- | --- |
| DAPI | Thermo Fisher Scientific | Cat # D1306 |
| Anti-Iba1 | Fujifilm Wako | Cat # 019-19741 |
| Anti-PAX6 | BioLegend | Cat # 901301 |
| Anti-GFAP | Santa Cruz Biotechnology Inc. | Cat # SC-33673 |
| Anti-Casapse-3 | Cell Signaling Technology | Cat # 9661S |
| Anti-Ki67 | Invitrogen | Cat # MA5-14520 |
| Anti-SATB2 | Abcam | Cat # AB51502 |
| Anti-CTIP2 | Abcam | Cat # AB 28448 |
| Anti-Mouse 555 | Thermo Fisher Scientific | Cat # A21422 |
| Anti-Rabbit 647 | Thermo Fisher Scientific | Cat # A21244 |
| APP | Millipore | MAB348 |
| AT8 (pTau) | Thermo Fisher | MN1020 |
| **Medication** |  |  |
| Cyclosporine (Sandimmune Injection) | Novartis | NDC 0078-0109-01 |
| Dexamethasone Sodium Phosphate | Mylan | NDC 67457-420-10 |
| Bupivacaine | Fresenius Kabi | NDC 63323-466-17 |
| Necrostatin-1 | Enzo Life Sciences | CAS # 4311-88-0 |
| Sylgard silicone elastomer kit - polydimethylsiloxane (PDMS) | Electron Microscopy Sciences | Cat # 24236-10 |
| Meloxicam (2 mg/kg) | Norbrook | NDC 79926-070-17 |
| Euthasol | Virbac | NDC 051311-050-01 |
| **Cell lines** |  |  |
| Human WT iPSC line "C1-2" derived from healthy fibroblasts (male) | ATCC fibroblasts (CRL-2097); (Wen et al., 2014) | C1-2 |
| **Chemical Reagents and Resources** |  |  |
| Matrigel | Corning | Cat # 354277 |
| mTESR Plus basal medium | StemCell Technologies | Cat # 100-1130 |
| ReLeSR | StemCell Technologies | Cat # 100-0483 |
| DMEM/F12 | Gibco | Cat # 11330032 |
| KOSR | Gibco | Cat # 10828028 |
| Penicillin-Streptomycin | Gibco | Cat # 15140122 |
| GlutaMAX | Gibco | Cat # 35050061 |
| MEM-NEAA | Gibco | Cat # 11140050 |
| β-mercaptoethanol | Gibco | Cat # 21985023 |
| N2 Supplement | Gibco | Cat # 17502048 |
| Neurobasal | Gibco | Cat # 21103049 |
| B27 Supplement | Gibco | Cat # 17504044 |
| Normal goat serum | Cell Signaling Technology | Cat # 5425S |
| Fluoromount-G | Thermo Fisher Scientific | Cat # 5018788 |
| **Small Molecules** |  |  |
| LDN193189 | StemCell Technologies | Cat # 72147 |
| SB431542 | StemCell Technologies | Cat # 72234 |
| Y-27632 Dihydrochloride | StemCell Technologies | Cat # 72304 |
| CHIR99021 | StemCell Technologies | Cat # 100-1042 |
| Ascorbic Acid | StemCell Technologies | Cat # 72132 |
| BDNF | Gibco | Cat # 450-02-50UG |
| GDNF | Gibco | Cat # 450-10-50UG |
| Dibutyryl-cAMP | StemCell Technologies | Cat # 100-0244 |
| **Equipment** |  |  |
| 6-well plates | CELLTREAT Scientific Products | Cat # CT-229105 |
| NanoZoomer S360 Digital slide scanner | Hamamatsu | Cat # SBIS0121E04 |
| Universal Mycoplasma<br>Detection Kit | ATCC | Cat # 30-1012K |
| **Software and Algorithms** |  |  |
| ImageJ (Fiji) | NIH | ImageJ<br>https://imagej.net/ij/docs/guide/146-2.html<br>- RRID:SCR_002285 |

## Acknowledgements

This work was supported by the National Institutes of Health (R01NS119472 to HIC) and the Department of Defense (W911NF2310276 to HIC, GP, and CCB).

Opinions, interpretations, and conclusions are those of the authors and are not necessarily endorsed by the National Institutes of Health or Department of Defense.

